# A decision-making tool to navigate through Extracellular Vesicle research and product development

**DOI:** 10.1101/2023.11.16.567368

**Authors:** Francesca Loria, Sabrina Picciotto, Giorgia Adamo, Andrea Zendrini, Samuele Raccosta, Mauro Manno, Paolo Bergese, Giovanna L. Liguori, Antonella Bongiovanni, Nataša Zarovni

**Author notes:** **Correspondence**: Francesca Loria and Nataša Zarovni, HansaBioMed Life Sciences Ltd, Tallinn, Estonia. and.

## Abstract

Due to their intercellular communication properties and their involvement in a wide range of biological processes, extracellular vesicles (EVs) are increasingly being studied and exploited for different applications. Nevertheless, their complex nature and heterogeneity, as well as the challenges related to their isolation, purification and characterization procedures, require cautious assessment of the quality and quantity parameters to monitor. This translates into a multitude of choices and putative solutions that lie in front of any EV researcher, in both research and translational environments, resembling a labyrinth with multiple paths to cross and, possibly, more than one exit. In this respect, decision-making tools might represent our modern Ariadne’s string to follow not to get lost or distracted along the journey, to choose the shorter and best-fit-to-source EV application(s) and *vice versa*. Here, we present the implementation of a multi-criteria EV decision-making grid (EV-DMG) as a novel, customizable, efficient and easy-to-use tool to support responsible EV research and innovation. By identifying and weighting key assessment criteria for comparing distinct EV-based preparations and/or processes, our EV-DMG may assist any EV community member in making informed, transparent and reproducible decisions regarding the EV sources and/or samples to be managed, as well as the most suitable production and/or analytical pipelines to be adopted for targeting a defined aim or application.

## 1. Introduction

Extracellular vesicles (EVs) are nanosized membranous particles secreted and uptaken by either eukaryotic or prokaryotic cells. By intrinsically conveying a multitude of biologically active components, including, yet not limited to, proteins, lipids, nucleic acids and metabolites, EVs have shown to mediate cell-to-cell and organism-to-organism communication at both intra-taxon and inter-taxa levels, thus governing the functionality of any organism, as well as its response to and/or impact on the surrounding environment [1,2].

The complex and heterogeneous nature of EVs underly both the great potential and the challenges relative to their exploitation as intrinsic, “all-in-one” biological platforms for exploring and addressing multiple biological issues in a comprehensive manner. Hence, lately and at a remarkable pace, the spotlight on EVs has entailed a thriving repertoire of hopes and expectations, leaning on the promise of EV-based technologies to revolutionize several domains of nanoscience and offer competitive clinical and commercial solutions within massive industrial landscapes (*e.g.*, healthcare, personal care and agrifood) [3]. In line with this, conspicuous public funding, alongside corporate and venture investments, has supported the prominent academic and industrial engagement into intensive Research and Development (R&D) initiatives, as well as first Product Development (PD) attempts.

Within the perimeter of such a compelling innovation space, preeminent techno-futurist visions of EV-based solutions have contributed to raise social acceptance on a wide range of EV application scenarios. However, the biological unpredictability and irreducibility characterizing the composition of EV source(s) and derived EVs preside over the enduring gap between current investments and the actual EV technology-regulatory readiness [3,4,5]. The shortage of reference materials, quality standards and standardized methodologies for EV R&D and PD still challenges our research ability to unravel the full spectrum of either inherent or “acquired” EV biocomponents (*e.g.*, EV corona that become associated with EVs during or later upon their production and extracellular release), as well as to recognize EV active moieties and mechanisms of action. Consequently, this poses a burden not only on understanding and resolving the true structure, biocomposition and physio-pathological role of EVs, but also on the ability to reproducibly and reliably manufacture EVs.

Currently, the EV bioprocessing panorama in either (basic and preclinical) research or clinical and/or industrial settings is mostly dominated by non-standardized, non-automated and “open” EV manufacturing systems [6,7,8,9,10,11,12,13]. On one hand, this enables keeping the pass with rapid innovation in the field, considering both known and unknown “unknowns” [14]. On the other hand, it ordinarily associates with the inevitable introduction of variability into EV-based products, resulting in inconsistencies in their structural identity (*e.g.*, EV subset collection), quality and safety, and accounting for the high discrepancies highlighted among distinct studies. By potentially and severely affecting data reliability and reproducibility, this overall lack of standardization and robustness exerts a strong, hostile impact on the EV landscape. It also challenges the accuracy by which scientists, nowadays, select the most appropriate applications to be tailored to any given EV source, as well as to formulate and dose EVs. On top of that, predominant EV bioprocessing procedures are still mostly research-compliant, characterized by considerable cost intensiveness and low-throughput performance, which are able to address, to date, only small-scale preclinical development and/or integration into clinical trials enduring to feature small patient cohorts (*e.g.*, 10-500 enrollments), even at advanced study phases (*i.e.*, phases 3-4) [15]. Proper support to EV manufacturing and analytical assessment at industrial scale, requiring extremely controlled standards of precision and limited space for errors, is still an open issue.

Working on EVs, therefore, constitutes a persisting “borderline” scenario where opportunities and risks coexist, inducing EV decision-makers to pursue their conquest of understanding, evidence and solutions while being continuously accompanied by the dynamic exposure to uncertainties related to what is not yet acquainted and/or may be not controlled [16]: unpredictability and irreducibility of EVs and related biological systems; lack of adequate methodological tools to unlock EV complexity; the potential emergence and/or impact of external risk factors and threats. In this light, to prompt the translation into effective, viable and marketable solutions, EV innovation needs to be a streamlined and a responsible one, where researchers themselves set the discovery and developmental stages by the early adoption of tailored, standardized, high-throughput, scalable and cost-effective working methodologies in a quality controlled and Good Manufacturing Practice (GMP)-oriented R&D and PD environment. Considering that “innovation is about change” [16], responsible Research and Innovation (RR&I), also in the EV field, should entail proper risk management beforehand and along the bioprocessing path. Especially when it comes to challenging and/or not well characterized study objects, as in the case of EVs, we ought to shape our innovation with a view to gaining a “dynamic capability” that may rapidly and duly “reframe” the R&I process in response to unpredictable and/or complex constraints, thus minimizing the emergence of unintended impacts and embedding cost intensiveness [16]. To reach this goal, EV R&D and its supporting activities related to quality, data and risk management should meet, mix and contaminate more and more. This could result in the production of interface tools that could be customized, tailored to research and researcher identity, which may truly support, and not cage, knowledge [11], while making it truly fruitful for the social wellbeing.

Hereby, we describe the application of a multi-criteria decision making (MCDM) matrix or grid to implement a customized, effective, easy-to-use, transferable and standard-oriented decision-making tool (DMT) model, specifically dedicated to EV features and related bioprocesses, which we named EV-DMG. In this manuscript, we showcase the adoption of our EV-DMG to streamline and improve in-process and/or end-product Quality Control (QC) of EVs derived from microalgae. We reveal its effectiveness in promoting application-tailored EV classification based on measurable quality scores, in accordance with pre-defined application-centric acceptance criteria, leading to minimization of erroneous and cost-intensive misuses. Furthermore, we describe the great potential of implementing our EV-DMG to: i) maximize EV bioprocess cost and turnaround time efficiency targeting the achievement of desired quality and quantity performances; ii) provide “real-time” traceable records of the whole EV PD cycle to be ultimately considered for final EV-based product regulatory dossier compilation. Overall, we spotlight the EV-DMG as a valid tool to steer complex decision-making throughout the entire EV value chain, from research to PD, with the ultimate goal of delivering responsible and sustainable EV R&I.

## 2. Comprehensive methodology for EV-DMG design and application

The workflow for the design of the EV-DMG is displayed in Figure 1. Overall, the process format comprises four main steps: (i) identification of “alternatives”, which represent the EV-based preparations and/or processes to compare for evaluation and decision-making; (ii) definition of EV-related alternative assessment criteria and relative importance weight; (iii) determination of EV-related alternative acceptance criteria and relative scoring system; (iv) implementation of the MCDM method to integrate EV-related alternative assessment into a Weighted Sum Model (WSM) matrix to perform alternatives’ ranking. The elements and concepts at the basis of our MCDM approach are elucidated in the following sections.

**Figure 1.**
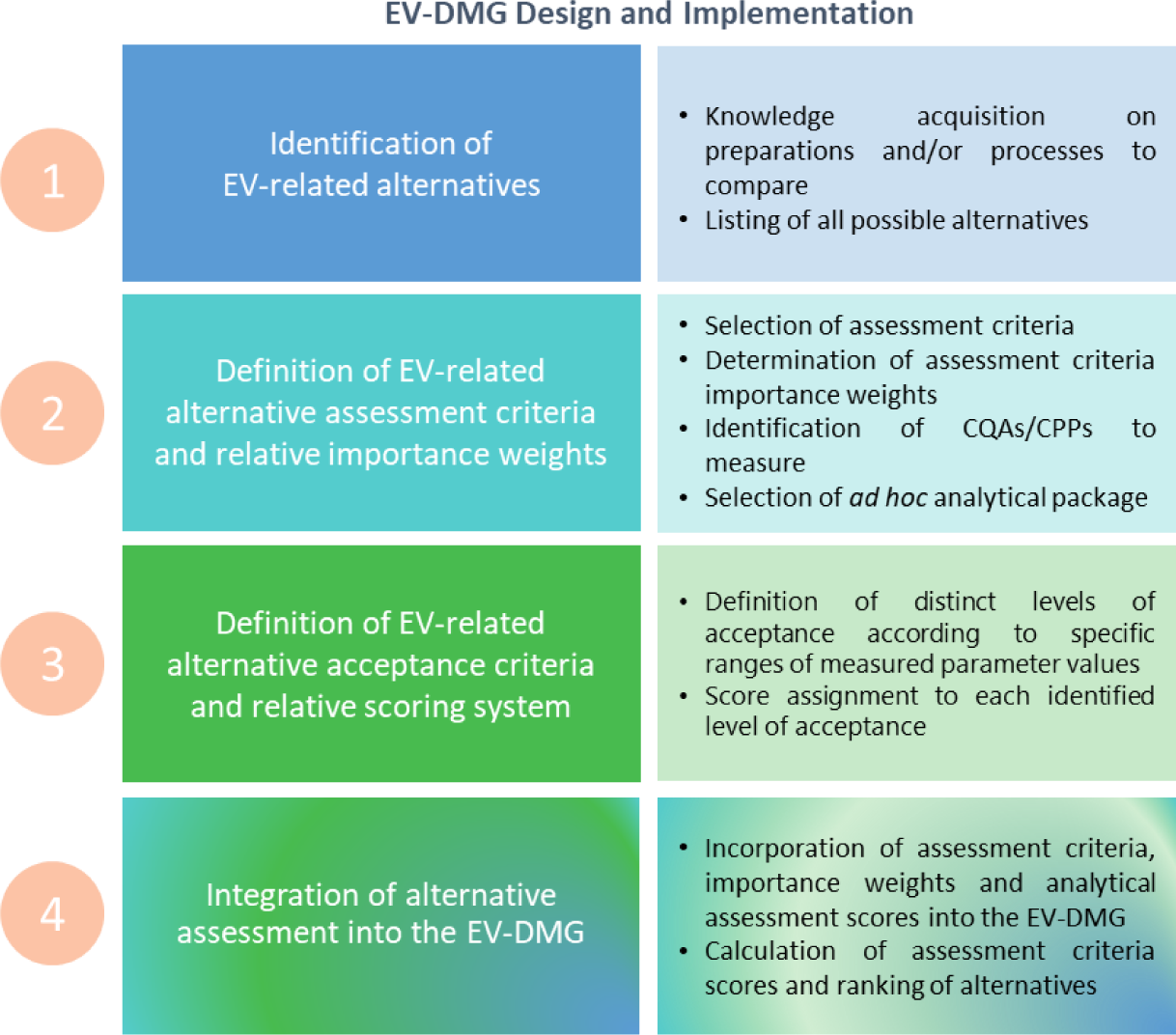
Proposed workflow for EV-DMG design and implementation.

### 2.1. Step 1 – Identification of EV-related alternatives

A MCDM approach generally targets the generation of preferences from alternative choices, options or conditions (*i.e.*, “alternatives”), which need to be compared, by considering and integrating more than one assessment criterion in the selection process. In any EV R&D settings, building an EV identikit towards specific purposes is conditioned by several fundamental decisions on critical factors that require to be clearly defined and assessed – either confirmed, potentially compared with alternative solutions or changed. These are mainly the following: **EV source; methods for EV production, handling, characterization and QC; EV applications**. The above enlisted and below described variables are primary, interconnected constituents of the comprehensive experimental setup featuring any EV R&D project, which drive the definition of the fundamental assessment criteria that should be established for designing, evaluating, improving and maximizing EV-based product and process performance along the way towards intended goals.

#### 2.1.1. EV source(s)

A research group or a study typically focuses on a particular EV source, whether being of human, animal, plant, bacterial or another origin. The raw, input material, which is provided or produced upstream the EV bioprocess, includes the EV-containing medium or matrix, as well as the potential excipients used to improve EV stability or extraction. The selection of the EV source drives the selection of the downstream methods to be adopted for both EV isolation/separation and characterization. Most importantly, the EV source is the major determinant for the potential, ultimate applications that may be hypothesized due to prior known functional features of the source. However, the effects and consequent possible uses of the EVs selected as study model might be completely novel and unexpected at the beginning of the research journey. EV sources have inherent variability that may result from the type of producer cell (*e.g.*, blood cells, stem cells, epithelial cells, endothelial cells, etc.), the strain of the producer cell (*e.g.*, human, plant, bacterial, etc.) or between different donors [*e.g.,* different donors of blood, Mesenchymal Stem Cells (MSCs) and bovine milk]. Although such variables may be subjected to a certain extent of standardization and control to minimize their impact on outcomes, at the same time, all these samples differ in complexity and contain different macromolecular contaminants or co-effectors, some of which might impose safety concerns. Finally, different EV sources come into play in different scales and offer different sustainability in terms of environmental footprint and economic viability. While these elements are not always priority criteria for fundamental researchers, they gain upmost importance in later translational phases versus productization and, according to RR&I principles, they should be considered in advance. Although this manuscript focuses largely on downstream EV bioprocessing, we appraise the importance of the upstream input material collection for promoting reproducibility, robustness and scalability of the whole bioprocess. In addition, we acknowledge that some key principles of decision-making, as well as the use of our EV-DMG itself, may be easily extended to also embrace the EV bioprocess upstream phase.

#### 2.1.2. EV isolation and purification method(s)

The selection of the methods to perform EV isolation and purification affects the quantity, quality, purity or enrichment of isolated EVs with respect to co-isolated molecules or particles. The appropriate isolation method will depend on the complexity and scale of the input material, as well as the needs for yield, purity and integrity of the extracted EVs. Some applications will call for high purity or even enrichment of specific EV subtypes (*e.g.*, if they are used as drug delivery vectors), whereas others would allow or even benefit from the presence of bioactive co-isolates (*e.g.*, if EVs are used as effectors in regenerative medicine). In any case, the yield and composition of EVs needs to be controlled and reproducible, while targeted function or feature must be preserved during the whole downstream processing. In a manufacturing system, yield is a critical parameter to be estimated for assessment of process performance, as outlining the percentage of non-defective produced items that have passed quality check for downstream processes and/or applications. For some EV types, such as microalgae-derived EVs (*i.e.*, nanoalgosomes) [17,18,19], for which we still lack clinical data that can allow for the accurate calculation of relevant scales needed to support (clinical) application, the scalability of the selected isolation and purification method is a preferential feature to consider for ensuring translatability. It is possible to do preliminary estimations based on the known precedent scenario for cell therapeutics, in particular initial data resulting from MSCs or other human cells that were the first ones to hit large scale clinical trials in humans. We will first have the estimate of an effective dose (per *in vitro* or *in vivo* experiment) to back-calculate the size of a “production lot” or even make first anticipation for in-human use planning. Of course, for a given desired effect, the dose is not just quantity, but also potency. Therefore, the yield requirements will differ considering the combination of EV source and indication for use. In general, downstream process (EV isolation and purification from the raw, input material) accounts for more than 50% of cost-of goods (COGs) for the whole bioprocess and this becomes a critical feature of EV manufacturing for widespread EV deployment, along scalability and efficacy [20]. In research phase, the issue of costs for both isolation and analytics can impact research pace and quality, with persistent inequalities across different labs and geographies. Therefore, the selection of the most suitable methods and protocols for performing EV isolation and purification would benefit from the use of a DMT, as our EV-DMG described below, to drive scientific and economic evaluations.

#### 2.1.3. EV sample handling and storage method(s)

Handling and storage procedures for both raw materials and harvested EVs may have a significant impact on EV quality and performance. Lack of proper evaluation on best or detrimental storage practice may increment variables in expected quantitative and qualitative gains. Costs associated with storage, packaging and shipment are currently assumed not to vary for different EV purification technologies or origins, but this is also something that needs to be proven. The impact and selection of storage protocols may be properly evaluated using our EV-DMG introduced below.

#### 2.1.4. Characterization method(s)

In the case of biotechnological/biological/chemical entities designed to be formulated for specific applications, the process of measuring requires the pre-identification of any physical, chemical, biological and/or microbiological properties associated with the desired product to be developed, conventionally defined as critical quality attributes (CQAs), which ought to be monitored and maintained within acceptance limits, ranges or distributions to ensure targeted product quality. It also demands the determination of the critical material attributes (CMAs) and critical process parameters (CPPs) that could affect targeted product’s CQAs. Measurements and data analytics necessarily underpin the development, control and validation of any (bio)process, including the EV-related one, being fundamental to make informed decisions and optimize operations. Measurable outputs give the key input for decision-making during (bio)process/product monitoring and benchmarking against references or alternatives. A large repertoire of methodologies is today available to address EV number, size, morphology, physiochemical and/or biochemical properties (Figure 2). The formulation of an analytical package that may fit the targeted scale (laboratory or industrial), sample type and EV application is both a result and a tool for evidence-based decision-making planning and decision-making. Currently, there is no possibility of relying upon a universal analytical package for all EV R&D or PD projects; instead, it requires to be *ad hoc* established, although many common methods will be recurrent and overlapping. The preferred analytical tools would ideally provide measurable readouts of the bioactive component(s) of the EVs of interest directly or indirectly contributing to their mechanism of action. Alternatively, in any case, it would be capable of screening and classifying the EV-containing samples at different stages, from raw to purified samples, thus enabling timely and accurate adjustment, as well as decrease in variability (*e.g.*, allowing us to choose the right donor).

**Figure 2.**
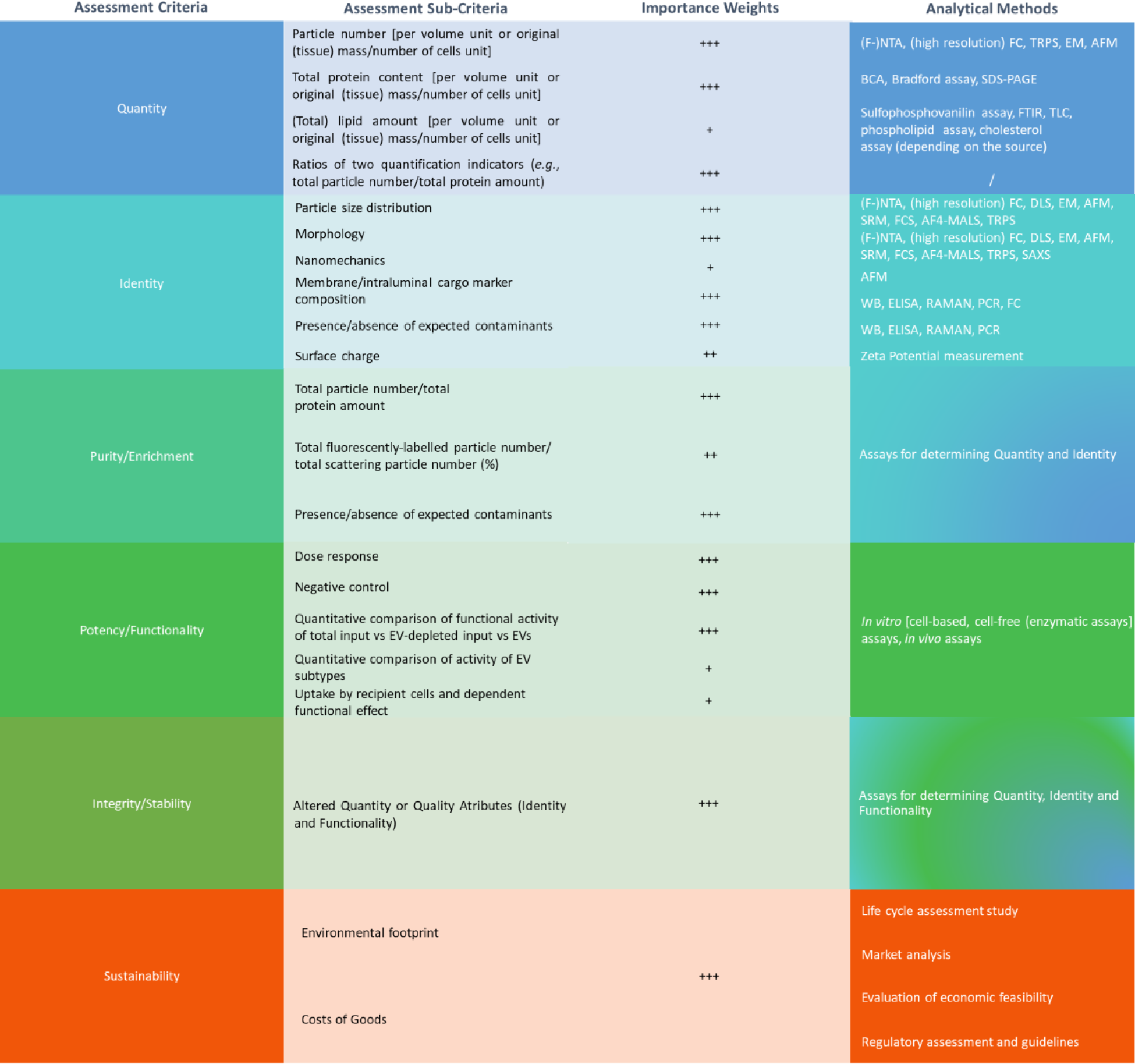
Setting the EV-DMG metrics for EV R&D – alignment of assessment criteria, sub-criteria, MISEV2018-adapted importance weights and analytical methods.

#### 2.1.5. EV applications

Evolutionarily speaking, the outstanding, comprehensive variety and complexity of EVs translate into a wide range of possible and appealing application scenarios. Focusing on one or a few uses is further complicated by the pleiotropic and multiplex nature of EVs, raising the possibility that single-sourced EVs might gain, and even monetize, high value in multiple ways. Nevertheless, the extended consideration and handling of EVs as a sort of magical bullets for any purpose, as well as the tendency to put everything under the same umbrella, do not benefit the progression of the single research, nor the whole field. The well-reasoned and evidence-informed choice of the best application to pursue would not limit innovation in research, but it would rather transform research into innovation that can truly translate into improvement of human condition.

### 2.2. Step 2 – Definition of EV-related alternative assessment criteria and relative importance weights

For the design of an EV-DMG based on the MCDM approach, a set of qualitative and quantitative EV-related assessment criteria need to be formulated to drive decision-making and the selection of the best “alternative(s)” under evaluation. Each assessment (sub-)criterion identifies and addresses a specific question by evaluating one or more inherent EV-related qualitative and quantitative attributes (*i.e.*, CQAs/CPPs), ultimately yielding clear and measurable assessment results. Figure 2 outlines the six major assessment criteria considered in R&D contexts to qualify EV-based products and/or processes, most of them adapted from the checklist reported in the “Minimal Information for Studies of EVs” (MISEV) 2018 guidelines [7] and recurrently used in EV studies. Each assessment criterion, identified as quantity, identity, purity/enrichment, potency/functionality, stability/integrity and sustainability, may be subdivided into diverse sub-criteria, which, in turn, may be addressed by several analytical methods. MISEV2018 proposed the following system for criteria scoring: +++ for mandatory criteria; ++ for mandatory criteria, if applicable to follow; + for encouraged criteria (Figure 2). We assumed this weighing system would be equivalent to numerically assigning an importance weight ranging from 3 to 1, indicating high, medium or low importance, respectively, as later applied in Figure 3.

**Figure 3.**
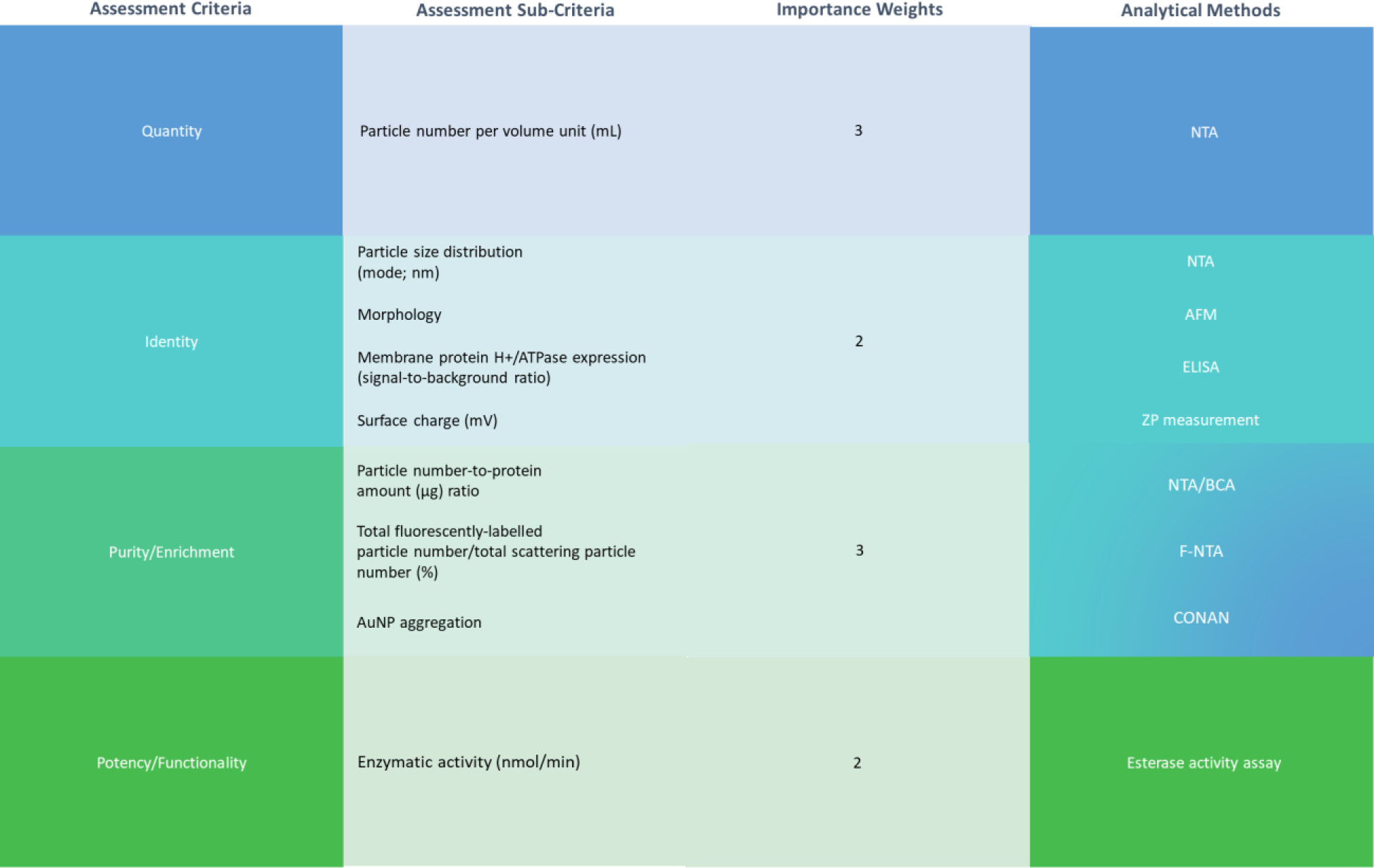
Setting the EV-DMG metrics for nanoalgosome QC – alignment of assessment criteria, sub-criteria, importance weights and analytical methods. Legend: AuNP = pristine gold nanoparticles.

### 2.3. Step 3 – Definition of EV-related alternative acceptance criteria and relative scoring system

Formatting an EV-DMG based on MCDM principles necessarily imposes the establishment of appropriate acceptance criteria defining the EV-based product/process requirements to be met to fulfil stakeholders’ expectations and needs for an intended purpose/application. EV acceptance criteria represent “numerical limits, ranges, or other suitable measures” [21] relative to the EV-based product’s and/or process’s CQAs and CPPs under assessment. These require to be empirically determined, for each appointed assessment criterion, by means of specific, pre-selected and pre-validated analytical procedures, and are ranked through a systematic scoring system, which reflects differential degree of requirement fulfillment. Therefore, in an EV R&D environment, the definition of acceptance criteria and corresponding scoring should be contextualized and justified based on empirical data demonstrating targeted EV-based product and/or process conformity to desired quality and functionality. The rationale and description of the acceptance criteria and respective scoring system conceived for showcasing the application of our EV-DMG are described in Section 3.3.

### 2.4. Step 4 – Integration of MCDM-based EV-related alternative assessment for ranking and application-centric classification

Among the plethora of strategies currently known and used to drive MCDM, the WSM represents one of the earliest and, possibly, the most commonly adopted method due to its easiness and effectiveness [22]. In our study, at this step, the relevant criteria selected for assessment of alternatives have been systematically incorporated into a WSM matrix to perform MCDM-based EV samples/conditions’ ranking or grading. The general construction of a WSM matrix satisfies the following mathematical expression [1,23]:

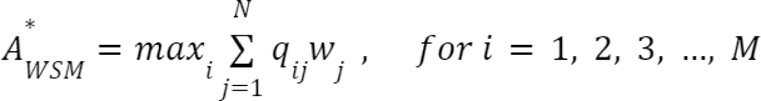

in which*: A*_WSM_* is the WSM of final score of the (best) alternative; *M* is the number of the alternatives to assess; *N* is the number of the decision criteria; *i* is a single alternative; *j* is a single criterion; *q_ij_* is the actual value relative to the i-th alternative in terms of the j-th criterion; *W_j_* is the importance weight of the j-th criterion. Simply put, upon generation of the WSM matrix, each decision or assessment criterion is given an analytical assessment score (*i.e.*, *q_ij_*), based on conformity to acceptance criteria, which is multiplied by its relative importance weight (*i.e.*, *W_j_*) to obtain an assessment criterion score. The sum of all assessment criterion scores provides a final, cumulative score or grade (*i.e.*, *A*_WSM_*) relative to each alternative evaluated, which leads to alternative positive selection for a specific downstream application or discarding. An illustrative example of the WSM matrix’s generation and exploitation is provided in the following subsections, showcasing our EV-DMG.

## 3. Illustrative example of EV-DMG design and application for nanoalgosome R&D

In compliance with the current recommendations by the EV community to integrate Quality principles and methods into EV R&D practices to foster rigor and standardization, it is not surprising that our EV-DMG found its first, immediate application in assisting quality management. QC is an essential aspect of quality management in EV R&D, as it ensures that the EV-based products and derived results of a study are consistent and effective. In EV R&D scenarios, QC should not be limited to review EV-based products against mere acceptance criteria for a single indication for use, but it should rather serve for matchmaking a given EV sample with the different uses it may have, driving our research to one of the possible directions it may take. In this perspective, a well-designed DMT may guide the appropriate QC actions, as well as support and test the unbiased hypotheses on the actual potential of a given EV sample for one or a few possible applications. In this manuscript, we exemplify the setting, adoption and utility of our EV-DMG to streamline QC assessment and grading of nanoalgosomes, chosen as model EVs. Its result addresses the consequent destination of our EV preparations to different uses within a multidisciplinary and explorative scenario.

### 3.1. Identification of nanoalgosome-related alternatives

Nanoalgosomes were extensively characterized and evaluated in several lines of research conducted either autonomously by our group or within multidisciplinary collaborative projects exploring their potential as intrinsic effectors or drug delivery vehicles in the field of biomedicine, therapeutics and cosmetics [17,18,19]. Such a wide spectrum of applications, if left open and pursued simultaneously without any prioritization, proved to be challenging for optimizing research efforts and resources. Our first, obvious conclusion was that the same input material was not of optimal quality and quantity for whatever application we could contemplate. Our EV-DMG is suitable for a QC evaluation that leads EV batches of different quality to be shunted towards the most appropriate application or to be eventually discarded, if not reaching the desired standards. Therefore, it may actively contribute to avoiding the risk of obtaining misleading results or no results at all. In this work, we showcase the application of the EV-DMG to compare and classify two independent nanoalgosome batches. As a case-study, we have selected the Biogenic Organotropic Wetsuits (BOW) project [funded within the frame of the Horizon 2020 Future and Emerging Technologies (FET) Proactive Program in 2021], which is ultimately aiming at exploiting nanoalgosomes as scaffolds for the production of (engineered) EV membrane-coated magnetic bead (evMBD) devices. In that, two primary applications of nanoalgosomes considered relevant for project Partners’ immediate needs were either use in engineering and/or *in vitro/in vivo* functional testing.

### 3.2. Definition of nanoalgosome assessment criteria and relative importance weights

The selection of the relevant set of EV assessment criteria to grade the nanoalgosome batches under assessment followed the principles previously described in Section 2.2. Our QC system was based upon a minimal but informative analytical package comprising a feasible number of assessment methods, all having the characteristic of being user-friendly, time-effective, mostly transferable and requiring low sample consumption. The panel of analytical tools assessed the following vesicular properties: EV i) **quantity** [determined and expressed in terms of particle concentration by nanoparticle tracking analysis (NTA)]; ii) **identity** [assessing scattering particle size distribution by NTA, morphology by atomic force microscopy (AFM), expression of nanoalgosome biomarkers (*i.e*. the plasma membrane protein H+/ATPase) by enzyme-linked immunosorbent assay (ELISA), surface charge by particle zeta potential measurement]; iii) **purity/enrichment** [labelling samples with an EV-specific lipidic dye and measuring the percentage of fluorescent particles over the total number of scattering particles by F-NTA; determining by the ratio between total particle number and total protein content; estimating purity from soluble co-isolated proteins by Colorimetric NANoplasmonic (CONAN) assay] [24,25]; iv) **potency/functionality** [determining EV esterase activity as proxy of EV bioactivity by a proprietary EV functional enzymatic activity assay] [26,27]. To specifically address the BOW project purposes mentioned in Section 3.1, and considering the importance weighting system described in Section 2.2, we provided an importance weight equivalent of 3 out of 3 (*i.e.*, high importance) to the EV assessment criteria quantity and purity/enrichment, whereas an importance weight of 2 out of 3 (*i.e.*, medium importance) was assigned to the EV assessment criteria identity and potency/functionality (Figure 3).

### 3.3. Definition of nanoalgosome acceptance criteria and determination of relative scoring

To assign to each nanoalgosome batch a specific quality score determining its suitability for final envisaged application(s), a set of acceptance criteria was empirically defined based on previously assessed nanoalgosome quality “standards”, which were produced and used at the laboratory scale for initial characterization studies. These studies were inclusive of a wide range of nanoalgosome attributes, without limiting or exhausting the nanoalgosome ultimate applications. The set of acceptance criteria relative to each EV assessment sub-criterion was complemented by a scoring system, assigning a specific analytical assessment score to the obtained measurement results. Any score provided to a given analytical assessment described to which extent the sample met a defined requirement for a specific assessment (sub-)criterion. Specifically for the work presented here, we defined a scoring system featuring an evaluation scale as follows: 0 = null; 1 = low; 2 = medium; 3 = high (Figure 4).

**Figure 4.**
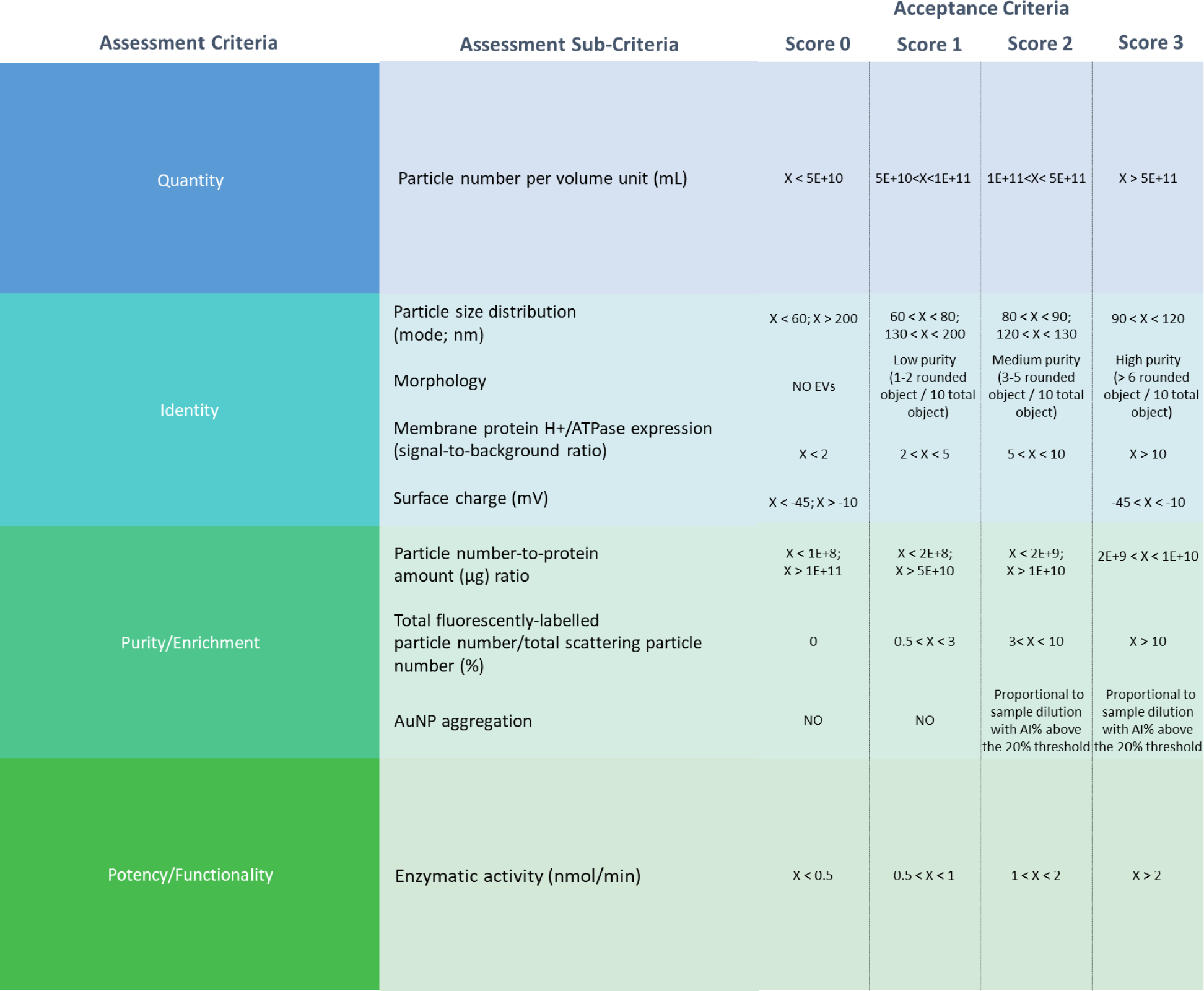
Setting the EV-DMG metrics for nanoalgosome QC – alignment of assessment criteria, sub-criteria and acceptance criteria with relative scoring. Legend: X indicates the measure of the corresponding parameter; AuNP = pristine gold nanoparticles; AI = aggregation index.

### 3.4. Integration of MCDM-based nanoalgosome assessment for ranking and application-centric classification

Figure 5 represents the EV-DMG compiled for the two batches of nanoalgosome taken under analysis, as described in Section 3.2 and in conformity with the acceptance criteria setup for nanoalgosome grading and classification, which are outlined in Section 3.3.

**Figure 5.**
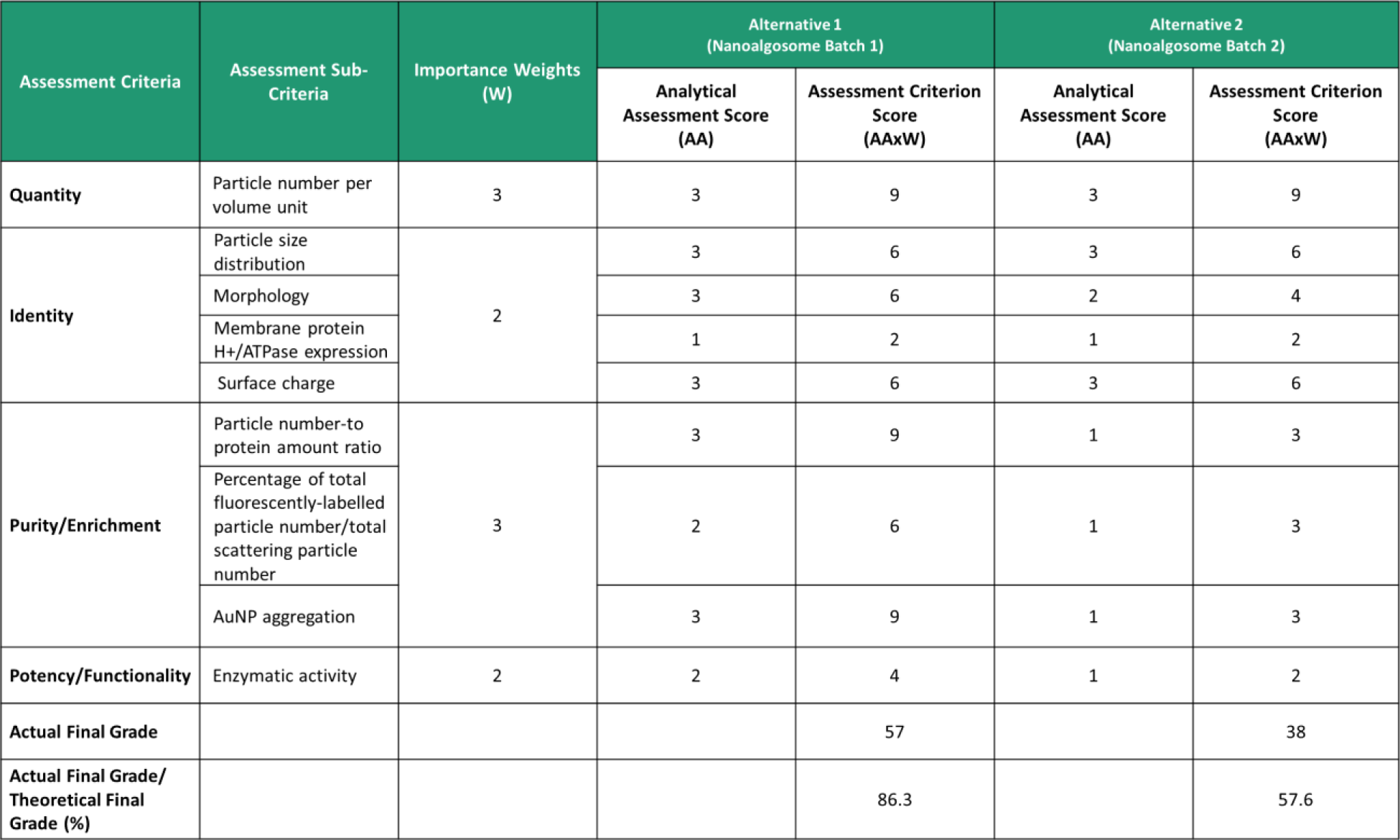
EV-DMG for nanoalgosome QC. Legend: AuNP = pristine gold nanoparticles.

The results obtained from the NTA analysis on nanoalgosome Batch 1 and Batch 2 (Figure S1) showed differences in terms of particle concentration (Batch 1 = 9E+11 ± 4.14E+10 particles/mL; Batch 2 = 8.6E+12 ± 4.56E+11 particles/mL), but a quite overlapping size distribution, peaked at about 100 nm. Not significant differences between the two batches were revealed neither by Z-potential measurements, indicating an analogous particle membrane surface charge, nor by H+-ATPase ELISA assay measuring the expression of a nanoalgosome-characteristic membrane protein (Figure S1). F-NTA on nanoalgosome labelled with a fluorescent lipophilic dye (di-8-ANEPPS), which emits fluorescence when anchored to the lipid bilayer, was used to perform purity evaluations by tracing fluorescent particles and identifying the presence of co-isolates. The ratio between fluorescent particles (Batch 1 = 3E+10 ± 2.6E+8 particles/mL; Batch 2 = 7.5E+10 ± 2.62E+9 particles/mL) and particles detected in scattering mode highlighted Batch 2 to display more co-isolated non-vesicular particles than Batch 1. In line with this observation, we estimated a different particle-to-protein ratio between the two batches (Batch1 1 = 4E+9 particles/µg proteins; Batch 2 = 7E+10 particles/µg proteins), further validated by CONAN assay, which confirmed the superior purity of Batch 1 from co-isolated proteins (Figure S1). Indeed, the particle-to-protein ratio of Batch 2 deviated from the proper ratio reported by Sverdlov (2012), stating that 1 µgr of EV preparation proteins corresponds to 2E+9 EVs [28]. In addition, AFM revealed in Batch 2 elongated objects potentially originating from microalgal cells or from the detachment of tangential flow filtration (TFF) cartridge fibers (Figure S1).

As shown in Table S1, the theoretical final grade was calculated by summing up the whole set of assessment criterion scores that could be obtained by multiplying the importance weight of each assessment criterion by the maximum analytical assessment score that could be potentially assigned to it, namely 3 out of 3. The observed discrepancies between the two batches translated into different analytical outputs on our EV-DMG. The sum of all assessment criteria scores, obtained by multiplying each analytical assessment score by its relative importance weight, resulted in an actual final grade equivalent to 57/66 for Batch 1 (86.3% of the theoretical final grade) and 38/66 for Batch 2 (57.6% of the theoretical final grade). As illustrated in Figure 6, depending on the actual final grade obtained, each nanoalgosome batch could be classified into one of three QC classification levels: level A, for nanoalgosome batches of good quality, which may be subjected to all uses, including engineering and functional studies, if confirmed sterile (actual final grade equal to 70-100% of the theoretical final grade); level B, for nanoalgosome batches of medium quality, which may be subjected to engineering and other uses, excluding functional testing (actual final grade equal to 30-69% of the theoretical final grade); level C, for nanoalgosome batches of bad quality, which require to be discarded or not to be used for critical analyses (actual final grade inferior than 30% of the theoretical final grade). Accordingly, the analysis and the use of our EV-DMG to streamline our nanoalgosome QC assessment highlighted, in a straightforward and objective manner, the suitability of Batch 1 to be exploited for further downstream analyses, including *in vivo* applications, and the potential for Batch 2 to be used only for engineering and non-critical analyses. In such a way, our group, as a nanoalgosome production site, could distribute the EV preparations to partner site in order to optimize the efforts and maximize the chance of success.

**Figure 6.**
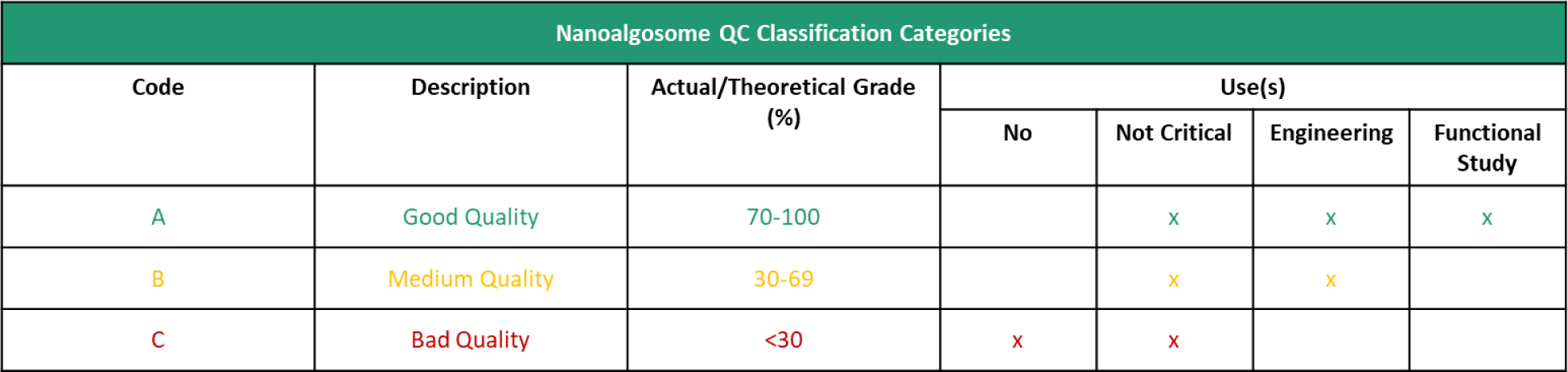
EV-DMG-based QC classification of nanoalgosomes.

## 4. Discussion

In this work, we focused our attention on the development and implementation of DMTs suitable for helping researchers, at any stage of their career and any level of competence in the EV field, to manage and address the multidimensionality of decision-making issues dominating and challenging the EV R&D and PD panorama. These may include either the accessibility to different EV sources, the variability in EV purity, composition and function within a single EV source, the plethora of EV isolation and characterization techniques available, as well as the wide spectrum of biologically and industrially relevant EV functions and applications contemplated and/or under investigation. Computational or mathematical DMTs have been historically adopted by decision makers to manage and mitigate the technical, logistical and economic risks associated with the complex set of factors and driving forces affecting system dynamics in numerous industrial manufacturing contexts (*i.e.*, from aerial and automotive vehicles to biotherapeutics) [29,30,31,32,33]. DMTs have also been applied in public research environments to help identify the most promising project directions, aims and activities [34,35]. In early 2019, the EV landscape has first embraced the notion of implementing DMTs for assisting research-grade and clinical-grade EV-based PD, when Ng and co-authors developed a novel computational DMT to identify combinations of technologies for upstream (human cell expansion) and downstream (EV isolation and purification) EV manufacturing to promote upscaling, while addressing minimization of COGs [20]. More recently, Picciotto et al. (2021) have introduced the adoption of a WSM-based DMT for addressing the selection of the microalgal sources best suited for microalgae-derived EV production [17]. Nevertheless, the comprehensive and straightforward decision-support framework to streamline and drive complex decision-making at each stage of EV-based research and PD, seamlessly integrating the two, ideally, is still missing. MCDM-based DMTs provide a valid, reliable and straightforward RR&I-tailored approach to make unbiased selections and classifications within complex sets of alternative options. Specifically, MCDM assessment minimizes room for evaluation error by regulating evaluations according to specific assessment criteria and accounting for their relative impact (or “importance weight”) on the decision-making process. As such, this strategy opens to opportunities for cost and time saving, increased data reliability and reproducibility, meanwhile enhancing project performance, simplifying project administrative work, as well as supporting conflict prevention and resolution among stakeholders [29,36,37]. It generally allows decision-making to take the lead over decision-control activities, thus easing the path towards the concretization of the desired outcomes.

The EV-DMG, described in this manuscript, proposes a user-friendly model of a MCDM matrix that may be applied to guide decision-making throughout the whole EV bioprocessing journey: from the “upstream” selection and management of starting materials to the “downstream” extraction, purification, QC and release of the intended EV-based products. In our case study, we illustrated the exploitation of the EV-DMG as a straightforward and effective QC-supporting tool, enabling EV QC ranking and classification tailored to envisaged applications and users. The nanoalgosome preparations taken under analysis shared the same microalgal strain input, culture protocol and purification method, yet giving two distinct batches we could choose between (our “alternative” decision objects). By means of a minimal, time-effective, easy-to-use and mostly transferable analytical package, we assessed both samples against specific quantity and quality parameters (assessment criteria), which were assigned an importance weight directly correlating with their significance for achieving our pre-fixed research project objectives, namely their engineering and, ultimately, their application as functional scaffolds for the manufacturing of evMBDs tested in *in vitro* and *in vivo* models. In relation to each assessment criterion and sub-criterion evaluated, enlisted in Figures 3, 4 and 5, a given analytical assessment score, ranging from 0 to 3, was provided to each sample, based on its conformity to the empirically pre-determined and pre-validated acceptance criteria, reported in Figure 5. Final integration of all assessment (sub-)criterion scores, resulting from the association of importance weights and analytical assessment scores, allowed us to allocate the two tested nanoalgosome batches into two distinct categories of use, despite being produced by an equivalent isolation and purification procedure: category A for batch 1, opening to all possibilities of envisaged applications, including engineering and bioactivity testing *in vitro* and/or *in vivo*; category B for batch 2, destinating only to functionalization.

We have selected nanoalgosomes as model EVs to validate our approach. Nevertheless, the proposed EV-DMG is suitable for application and extension to any other EVs. As we initiated our work with nanoalgosomes at a very early, research-grade EV bioprocess stage, we duly assumed that good practice would entail the selection of the proper assessment criteria to adopt in our MCDM workflow in compliance with internationally accepted guidelines. The position statement reported as MISEV, first published in 2014 and lastly updated in 2018, together with other excellent initiatives of the scientific community (*i.e*., EV-Track), has been established as an EV community-agreed framework providing guidelines on the proper procedural steps to follow and the common metrics to use to perform good EV research and reporting on specific EV-associated functionalities [7,38,39]. This was conceived to address rigor and standardization issues, fostering data reusability, reproducibility, and comparison [5,11,12,40]. MISEV2018 piloted the attempt to assign specific importance weights to a comprehensive set of methodological criteria, enlisted in the checklist contained in the guideline document, addressing the EV characteristics that have been mainly contemplated by EV researchers, and considered a key for conducting EV basic science and first PD efforts (*i.e*. quantity, identity, purity/enrichment, potency/functionality and stability/integrity). It is noteworthy to specify that MISEV2018 did not diversify the importance of each methodological criterion according to chosen EV sources, study scopes or envisaged EV-based application(s). Rather, it averaged the broad state of opinion, gathered up to that time, on the general importance of assessing given EV attributes in specific methodological settings for generating reliable and reproducible data, thus proposing a “one scoring fits all or most”. Although the design of our EV-DMG was executed taking into consideration MISEV2018’s recommendations and importance weighing system, we realized that the selection of the essential EV-related procedural criteria to follow, the assignment of their relative importance weights, the correlated analytical package to adopt for addressing specific EV production, handling, characterization and formulation objectives were likely to be done empirically, in a case-by-case manner, with a view to spotting the universal weighing patterns and requirements (or combinations thereof) related to key decisions to take (enlisted in Section 2.1). Although it is well recognized that, at least in premature research phases, it would be ideal and wise to include as many EV assessment criteria as possible into a research-grade EV-DMG [41], it is explicit that their choice and technical application would still depend on the specific logistic and study context. Most importantly, their impact on our go-no-go decisions would vary depending on the particular research objectives to address. The same criterion may be considered more critical for certain research purposes, while less important for others. For instance, it is more likely that preserving EV integrity and potency/functionality is more critical for researchers intending to test and propose their model EVs as intrinsic effectors, than for those that are going to engineer them with heterologous moieties or cargos. Conversely, quantity is likely to be critical for dosing EVs in general and might be universally weighed as a highly impactful parameter. On the other hand, high purity and the absence of any other non-vesicular contaminants may have a very different importance in distinct applications. Another common example is given by studies aimed at identifying novel EV-based biomarkers, in which the assessment criteria and sub-criteria considered most crucial to track would include EV sample purity, the specificity of the EV isolation methods, the sensitivity and reproducibility of biomarker detection assays, as well as the validation of biomarker candidates using appropriate statistical methods.

MISEV2018 principles have been recognized as an excellent landmark also by early industrial EV innovators. Nevertheless, what has emerged from diverse position documents that have been recently published on the matter, especially focusing on the development of EV-based products as next-generation cell-free-based therapeutics, is that requirements more compliant to clinical and industrial settings are demanded, necessarily imposing due considerations about the assessment (sub-)criteria and relative importance weights to be selected [41,42] when designing and building a clinical- or industrial-grade EV-DMG (Figure 7). Concerning purity assurance, for instance, if the use of the main MISEV2018 sub-criteria to observe for its characterization receives wide consensus (*e.g.*, ratio of two quantification figures, such as particle number and protein content; sample examination for the presence or absence of expected contaminants, etc.), the requirement of high purity at the expense of potency should be revisited. Having recognized that a completely pure EV preparation is unlikely to be produced, as well as being aware of the potential association of some “impurities” (perhaps, associated with the EV “protein corona”) with desired sample biological activity, the clinics-driven EV community is currently recommending not to insist on stringent purification procedures that would only result in loss of sample therapeutic activity, yet to define the most critical contaminants and propose duly adjusted acceptance criteria, ensuring minimization of inter-batch variability [41]. Upon the construction of an EV-DMG, this could be translated into the reduction of the importance weight relative to purity estimation (for example, importance weight of 2/3 in place of 3/3) or into the delineation of alternative acceptance criteria and relative scoring systems. On the other hand, if bioactivity evaluation is not a “must” in a research-grade scenario, depending on the investigational purposes, fulfilling specific functionality requirements represents a very critical aspect when developing products for in-human use. This, again, could turn the variable importance weight attributed to the assessment of EV functionality in research settings into a definite importance weight equal to 3/3 into an EV-DMG to be used in a clinical environment (*e.g.*, therapeutics, vaccines and drug delivery). Importantly, considerations about microbiological contamination and economic sustainability for EV large-scale production are often overlooked at early-stage EV bioprocessing stages. Sterility is a mandatory feature of any medicinal product, which requires to be met or to be managed since premature research phases for minimizing safety concerns at early-phase clinical trials, as well as developmental setbacks [43,44,45]. Therefore, microbiological security, with a high, if not the highest, importance weight score, should be added to the panel of EV assessment criteria to be incorporated into an EV-DMG driving decision-making throughout the whole value chain of EV-based therapeutics. Furthermore, there is a growing call for more pragmatic PD approaches, in agreement with existing regulatory guidelines, consisting in selecting not only the most effective, but also the most feasible methodologies and procedures to assess the most relevant in-process/end-product EV-based product parameters (*i.e.*, CQAs, CMAs, CPPs). In this, the primary goal is meeting economic sustainability requirements, imperative at advanced bioprocess developmental and manufacturing stages of products for massive use and consumption, such as therapeutics for widespread diseases, consumer care products or agrochemicals [42]. Accordingly, EV assessment (sub-)criteria related to costs and time management should be considered for integral incorporation into any MCDM-based tool guiding decision-making especially towards translational applications. *Inter alia*, adopting MCDM-based tools to drive, *ex ante*, the allocation of resources into fair and best suited categories of use may enable us to make financial trade-offs and prevent cost-intensive misuses.

**Figure 7.**
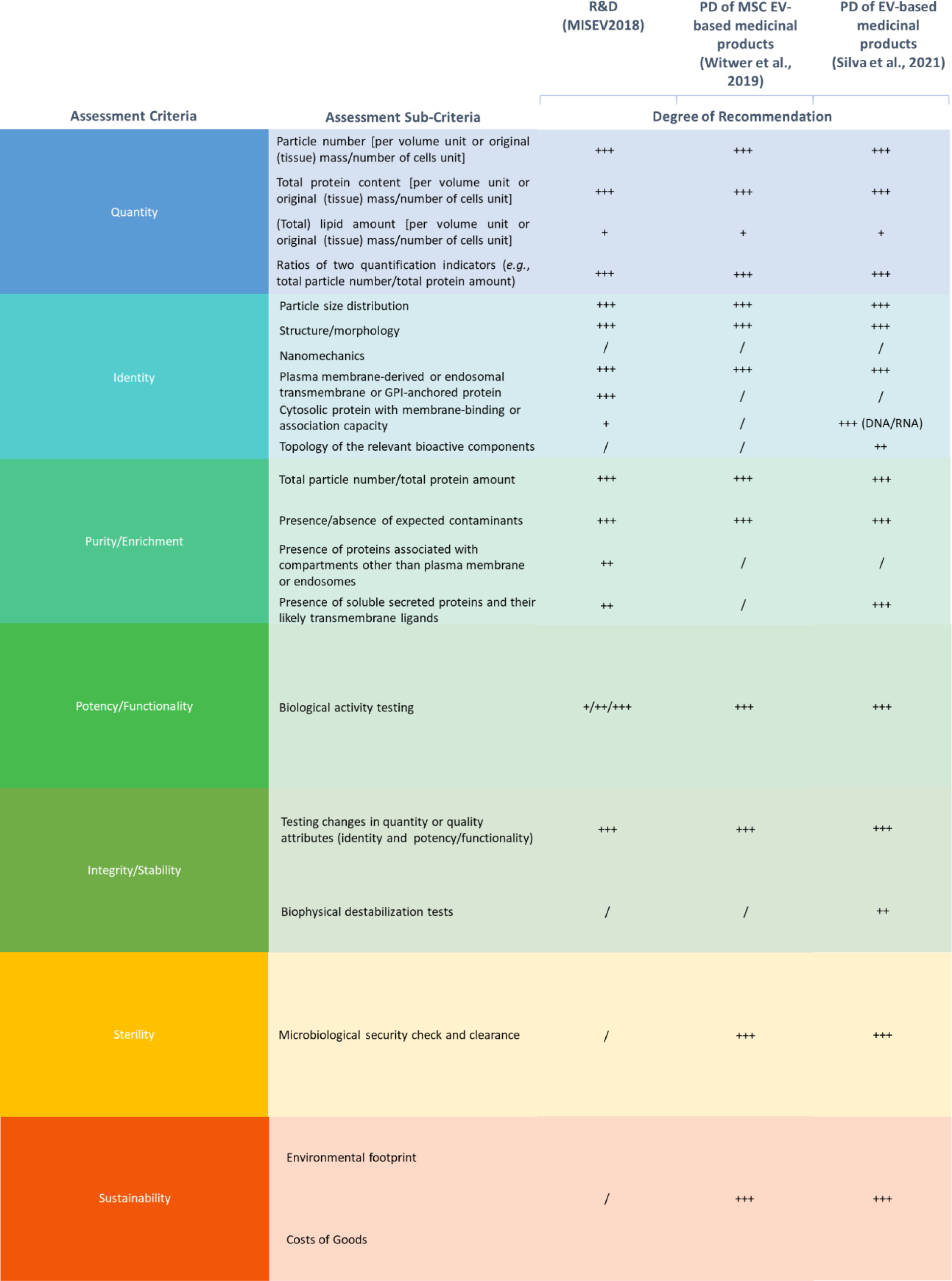
Setting the EV-DMG metrics for EV PD – alignment of assessment criteria, sub-criteria and importance weights deducted from MISEV2018 guidelines’ checklist, Silva, et al. (2021) [41] and Witwer et al. (2019) [42].

We have fully included all these considerations in the translation of nanoalgosomes as pure research objects into key elements of structured pipelines for delivery of novel formulations in cosmetics and drug delivery. Implementation of hereby described EV-DMG templates enables us to track back and forth correlations between obtained final grades and frontend decisions made regarding the culture conditions and substrates, as well as to reconsider the architecture of upstream and downstream manufacturing. On the other hand, we are able to trace the EV assessment criteria all the way to the desired performance of our nanoalgosome preparations in particular indications for use, link them directly to regulatory criteria and, thus, be informed on feasibility of our dosing and pricing strategies. This helps us, as any EV developer, to spot the unique advantages and “selling points” of nanoalgosomes as fit-to-solutions (*e.g.*, sustainability, stability, scalability, intrinsic activity) and enables us to iterate on our choices, tailor our scales and designs, while not limiting our capacity to further innovate and spin out novel nanoalgosome capabilities into innovative productization areas.

The integration of MCDM-based DMT systems, as our EV-DMG, into routine EV bioprocessing practices envisaging regulatory-compliant market authorization may arise as a tactical move to dynamically frame PD and manufacturing with early reference to late submission of product Common Technical Document (CTD) or product dossier. It could be aligned and merged into the Target Product Profile (TPP) of the prospective product to be developed, which, technically, represents a strategical and evolving document whose formulation is recommended, since product design phase, to guide benchmarking, provide a comprehensive recapitulation of the targeted and/or formulated product’s attributes and, overall, improve regulatory dialog for more efficient regulatory review and approval times [46]. Importantly, diverse National Regulatory Authorities (NRAs), including the European Medicines Agency (EMA) and the United States Food and Drug Administration (U.S. FDA), have been increasingly exploiting Multi-Criteria Decision Analysis (MCDA) to regulate approval decisions to optimize trade-offs between risks and benefits [47]. This suggests that the adoption of MCDM-based DMTs, as our EV-DMG, to support the performance of research and industrial settings targeting clinical validation and acceptance by health authorities could upgrade the communication and understanding between innovators and regulatory bodies about product value propositions, thus ultimately enhancing the likelihood of shortening product’s development cycle and market approval. Moreover, the EV-DMG templates that we propose may be easily handled by common software and associated scores, as numerical outcomes and indicators of evaluation, could be turned into algorithms providing an interface for unbiased and clear gatekeeping decisions.

## 5. Conclusions

Our work emphasizes the significance of fully integrating pivotal principles of RR&I into routine EV R&I practices by early conceiving, developing and/or adopting multi-dimensional methodological strategies that could foster “weighted” decisions for promoting robust, sustainable, ethically acceptable and socially desirable outcomes of EV science and innovation [16,48,49]. Hereby, we describe the adoption of a DMT to promote quantifiable, objective and auditable MCDA at each phase of the EV bioprocessing route. By showcasing our EV-DMG as a support tool for assisting QC of microalgae-derived EVs, we highlight the capability to perform accountable and impartial *a posteriori* qualification of in-process and/or end products/processes. This is achieved through the integration of systematic, regulatory-compliant MCDA with the expertise about EV input material’s and output product’s properties, along with the empirically accumulated awareness about targeted EV-based product requirements for specific downstream applications. Furthermore, as the relevance of a given assessment criterion may vary depending on the specific context of evaluation, and its relative importance weight may change accordingly, we also show the potential of this system to *a priori* tailor our resources and products towards *ad hoc* applications, ultimately maximizing overall EV bioprocess cost and turnaround time efficiency. This is relevant not only to sustain and boost the performance of R&D activities within the frame of multicentric and multidisciplinary projects, but also to enhance the prospects of successfully translating R&D efforts into regulatory-compliant marketable solutions. Finally, by providing “real-time”, traceable records of the whole EV PD cycle to be ultimately considered for final EV-based product regulatory dossier compilation, we further spotlight the promise hold by the EV-DMG to be leveraged as a valuable regulatory resource to promote well-structured regulatory communication and, consequently, more efficient, faster-paced review and approval times by NRAs.

## 6. CRediT authorship contribution statement

**Francesca Loria**: Conceptualization; Methodology; Investigation; Validation; Data Curation; Visualization; Writing– Original Draft Preparation; Writing–Review & Editing. **Sabrina Picciotto**: Conceptualization; Methodology; Investigation; Validation; Data Curation; Visualization; Writing–Original Draft Preparation; Writing–Review & Editing. **Giorgia Adamo**: Conceptualization; Methodology; Investigation; Validation; Data Curation; Writing–Original Draft Preparation; Writing–Review & Editing. **Andrea Zendrini**: Investigation; Validation; Writing–Review & Editing; **Samuele Raccosta**: Investigation; Validation. **Mauro Manno:** Resources; Investigation; Validation**. Paolo Bergese**: Funding Acquisition; Project Administration; Resources; Writing–Review & Editing. **Giovanna Liguori**: Conceptualization; Methodology; Writing–Review & Editing. **Antonella Bongiovanni**: Conceptualization; Methodology; Resources; Writing–Review & Editing; Supervision. **Nataša Zarovni**: Conceptualization; Methodology; Resources; Writing–Original Draft Preparation; Writing–Review & Editing; Supervision. Nataša Zarovni and Antonella Bongiovanni equally contributed to this work.

## 7. Acknowledgements

This work was funded by the Horizon 2020 Framework Programmes under the grant FETPROACT-952183 BOW.

## References

1. Raposo G, Stoorvogel W. Extracellular vesicles: exosomes, microvesicles, and friends. Journal of Cell Biology. 2013 Feb 18;200(4):373–83.

2. Yáñez-Mó M, Siljander PR, Andreu Z, Bedina Zavec A, Borràs FE, Buzas EI, Buzas K, Casal E, Cappello F, Carvalho J, Colás E. Biological properties of extracellular vesicles and their physiological functions. Journal of extracellular vesicles. 2015 Jan 1;4(1):27066.

3. Zarovni N, Loria F, Zenatelli R, Mladenovic D, Paolini L, Adamo G, Radeghieri A, Bongiovanni A, Bergese P. Standardization and commercialization of extracellular vesicles. InExtracellular Vesicles 2021 Oct 20 (pp. 303-335).

4. Bruno I, Lobo G, Covino BV, Donarelli A, Marchetti V, Panni AS, Molinari F. Technology readiness revisited: a proposal for extending the scope of impact assessment of European public services. InICEGOV 2020 Sep 23 (pp. 369-380).

5. Nieuwland R, Falcón-Pérez JM, Théry C, Witwer KW. Rigor and standardization of extracellular vesicle research: Paving the road towards robustness. Journal of Extracellular Vesicles. 2020 Dec;10(2).

6. Lötvall J, Hill AF, Hochberg F, Buzás EI, Di Vizio D, Gardiner C, Gho YS, Kurochkin IV, Mathivanan S, Quesenberry P, Sahoo S. Minimal experimental requirements for definition of extracellular vesicles and their functions: a position statement from the International Society for Extracellular Vesicles. Journal of extracellular vesicles. 2014 Jan 1;3(1):26913.

7. Théry C, Witwer KW, Aikawa E, Alcaraz MJ, Anderson JD, Andriantsitohaina R, Antoniou A, Arab T, Archer F, Atkin-Smith GK, Ayre DC. Minimal information for studies of extracellular vesicles 2018 (MISEV2018): a position statement of the International Society for Extracellular Vesicles and update of the MISEV2014 guidelines. Journal of extracellular vesicles. 2018 Dec;7(1):1535750.

8. Clayton A, Boilard E, Buzas EI, Cheng L, Falcón-Perez JM, Gardiner C, Gustafson D, Gualerzi A, Hendrix A, Hoffman A, Jones J. Considerations towards a roadmap for collection, handling and storage of blood extracellular vesicles. Journal of extracellular vesicles. 2019 Dec 1;8(1):1647027.

9. Lener T, Gimona M, Aigner L, Börger V, Buzas E, Camussi G, Chaput N, Chatterjee D, Court FA, Portillo HA, O’Driscoll L. Applying extracellular vesicles based therapeutics in clinical trials–an ISEV position paper. Journal of extracellular vesicles. 2015 Jan 1;4(1):30087.

10. Paolini L, Monguió-Tortajada M, Costa M, Antenucci F, Barilani M, Clos-Sansalvador M, Andrade AC, Driedonks TA, Giancaterino S, Kronstadt SM, Mizenko RR. Large-scale production of extracellular vesicles: Report on the “massivEVs” ISEV workshop.

11. Liguori GL, Kisslinger A. Standardization and reproducibility in EV research: the support of a Quality Management System. InAdvances in Biomembranes and Lipid Self-Assembly 2021 Jan 1 (Vol. 33, pp. 175-206). Academic Press.

12. Liguori GL, Kisslinger A. Quality management tools on the stage: old but new allies for rigor and standardization of extracellular vesicle studies. Frontiers in Bioengineering and Biotechnology. 2022;10.

13. Agrahari V, Agrahari V, Burnouf PA, Chew CH, Burnouf T. Extracellular microvesicles as new industrial therapeutic frontiers. Trends in Biotechnology. 2019 Jul 1;37(7):707–29.

14. Margolis L, Sadovsky Y. The biology of extracellular vesicles: The known unknowns. PLoS biology. 2019 Jul 18;17(7):e3000363.

15. clinicaltrials.gov; key search terms: exosomes and extracellular vesicles.

16. Owen RJ, Bessant JR, Heintz M, editors. Responsible innovation. Chichester: Wiley; 2013.

17. Picciotto S, Barone ME, Fierli D, Aranyos A, Adamo G, Božič D, Romancino DP, Stanly C, Parkes R, Morsbach S, Raccosta S. Isolation of extracellular vesicles from microalgae: Towards the production of sustainable and natural nanocarriers of bioactive compounds. Biomaterials Science. 2021;9(8):2917–30.

18. Adamo G, Fierli D, Romancino DP, Picciotto S, Barone ME, Aranyos A, Božič D, Morsbach S, Raccosta S, Stanly C, Paganini C. Nanoalgosomes: Introducing extracellular vesicles produced by microalgae. Journal of extracellular vesicles. 2021 Apr;10(6):e12081.

19. Paterna A, Rao E, Adamo G, Raccosta S, Picciotto S, Romancino D, Noto R, Touzet N, Bongiovanni A, Manno M. Isolation of extracellular vesicles from microalgae: a renewable and scalable bioprocess. Frontiers in Bioengineering and Biotechnology. 2022 Mar 14;10:836747.

20. Ng KS, Smith JA, McAteer MP, Mead BE, Ware J, Jackson FO, Carter A, Ferreira L, Bure K, Rowley JA, Reeve B. Bioprocess decision support tool for scalable manufacture of extracellular vesicles. Biotechnology and bioengineering. 2019 Feb;116(2):307–19.

21. Elder D. ICH Q6A Specifications: test procedures and acceptance criteria for new drug substances and new drug products: chemical substances. ICH Quality Guidelines: An Implementation Guide. 2017 Sep 27:433–66.

22. Triantaphyllou E, Shu B, Sanchez SN, Ray T. Multi-criteria decision making: an operations research approach. Encyclopedia of electrical and electronics engineering. 1998 Feb;15(1998):175-86.

23. Fishburn PC. Additive utilities with incomplete product sets: Application to priorities and assignments. Operations Research. 1967 Jun;15(3):537–42.

24. Zendrini A, Paolini L, Busatto S, Radeghieri A, Romano M, Wauben MH, Van Herwijnen MJ, Nejsum P, Borup A, Ridolfi A, Montis C. Augmented COlorimetric NANoplasmonic (CONAN) method for grading purity and determine concentration of EV microliter volume solutions. Frontiers in bioengineering and biotechnology. 2020 Feb 12;7:452.

25. Maiolo D, Paolini L, Di Noto G, Zendrini A, Berti D, Bergese P, Ricotta D. Colorimetric nanoplasmonic assay to determine purity and titrate extracellular vesicles. Analytical chemistry. 2015 Apr 21;87(8):4168–76.

26. 10.1101/2023.10.24.563745

27. Patent: No. 102023000004503

28. Sverdlov ED. Amedeo Avogadro’s cry: what is 1 µg of exosomes?. Bioessays. 2012 Oct;34(10):873-5.

29. Rao RV. Decision making in the manufacturing environment: using graph theory and fuzzy multiple attribute decision making methods. London: Springer; 2007 Jun 6.

30. Rekhi R, Smith JA, Arshad Z, Roberts M, Bountra C, Bingham I, Brindley DA. Decision-support tools for monoclonal antibody and cell therapy bioprocessing: current landscape and development opportunities. BioProcess Int. 2015 Jul 5;13(11).

31. Chhatre S, Francis R, O’Donovan K, Titchener-Hooker NJ, Newcombe AR, Keshavarz-Moore E. A decision-support model for evaluating changes in biopharmaceutical manufacturing processes. Bioprocess and biosystems engineering. 2007 Jan;30:1–1.

32. Tan TE, Peh GS, George BL, Cajucom-Uy HY, Dong D, Finkelstein EA, Mehta JS. A cost-minimization analysis of tissue-engineered constructs for corneal endothelial transplantation. PLoS One. 2014 Jun 20;9(6):e100563.

33. Sutton RT, Pincock D, Baumgart DC, Sadowski DC, Fedorak RN, Kroeker KI. An overview of clinical decision support systems: benefits, risks, and strategies for success. NPJ digital medicine. 2020 Feb 6;3(1):17.

34. Bongiovanni A, Colotti G, Liguori GL, Di Carlo M, Digilio FA, Lacerra G, Mascia A, Cirafici AM, Barra A, Lanati A, Kisslinger A. Applying Quality and Project Management methodologies in biomedical research laboratories: a public research network’s case study. Accreditation and Quality Assurance. 2015 Jun;20:203–13.

35. Digilio FA, Lanati A, Bongiovanni A, Mascia A, Di Carlo M, Barra A, Cirafici AM, Colotti G, Kisslinger A, Lacerra G, Liguori GL. Quality-based model for Life Sciences research guidelines. Accreditation and Quality Assurance. 2016 Jun;21:221–30.

36. Taherdoost H, Madanchian M. Multi-criteria decision making (MCDM) methods and concepts. Encyclopedia. 2023 Jan 9;3(1):77–87.

37. Zarghami M, Szidarovszky F, Zarghami M, Szidarovszky F. Introduction to multicriteria decision analysis. Multicriteria Analysis: Applications to Water and Environment Management. 2011:1–2.

38. Van Deun J, Mestdagh P, Agostinis P, Akay Ö, Anand S, Anckaert J, Martinez ZA, Baetens T, Beghein E, Bertier L, Berx G. EV-TRACK: transparent reporting and centralizing knowledge in extracellular vesicle research. Nature methods. 2017 Mar;14(3):228–32.

39. Hendrix A, Lippens L, Pinheiro C, Théry C, Martin-Jaular L, Lötvall J, Lässer C, Hill AF, Witwer KW. Extracellular vesicle analysis. Nature Reviews Methods Primers. 2023 Jul 27;3(1):56.

40. Hollmann S, Regierer B, D’Elia D, Kisslinger A, Liguori GL. Toward the definition of common strategies for improving reproducibility, standardization, management, and overall impact of academic research. InAdvances in Biomembranes and Lipid Self-Assembly 2022 Jan 1 (Vol. 35, pp. 1-24). Academic Press.

41. Silva AK, Morille M, Piffoux M, Arumugam S, Mauduit P, Larghero J, Bianchi A, Aubertin K, Blanc-Brude O, Noël D, Velot E. Development of extracellular vesicle-based medicinal products: A position paper of the group “Extracellular Vesicle translatiOn to clinicaL perspectiVEs–EVOLVE France”. Advanced Drug Delivery Reviews. 2021 Dec 1;179:114001.

42. Witwer KW, Van Balkom BW, Bruno S, Choo A, Dominici M, Gimona M, Hill AF, De Kleijn D, Koh M, Lai RC, Mitsialis SA. Defining mesenchymal stromal cell (MSC)-derived small extracellular vesicles for therapeutic applications. Journal of extracellular vesicles. 2019 Dec 1;8(1):1609206.

43. European Medicines Agency. ICH guideline Q8 (R2) on pharmaceutical development.

44. Guideline IH. Development and manufacture of drug substances (chemical entities and biotechnological/biological entities) Q11. London: European medicines agency. 2011 May.

45. Committee for Medicinal Products for Human Use. Guideline on the requirements for quality documentation concerning biological investigational medicinal products in clinical trials. London, UK: European Medicines Agency. 2012.

46. Breder CD, Du W, Tyndall A. What’s the regulatory value of a target product profile?. Trends in Biotechnology. 2017 Jul 1;35(7):576–9.

47. Chisholm O, Sharry P, Phillips L. Multi-criteria decision analysis for benefit-risk analysis by national regulatory authorities. Frontiers in medicine. 2022 Jan 12;8:820335.

48. Stilgoe J, Owen R, Macnaghten P. Developing a framework for responsible innovation. InThe Ethics of Nanotechnology, Geoengineering, and Clean Energy 2020 Jul 26 (pp. 347-359). Routledge.

49. Silva HP, Lefebvre AA, Oliveira RR, Lehoux P. Fostering Responsible Innovation in Health: an evidenceinformed assessment tool for innovation stakeholders. International journal of health policy and management. 2021 Apr;10(4):181.

50. Radeghieri A, Alacqua S, Zendrini A, Previcini V, Todaro F, Martini G, Ricotta D, Bergese P. Active antithrombin glycoforms are selectively physiosorbed on plasma extracellular vesicles. Journal of Extracellular Biology. 2022 Sep;1(9):e57.

