## Supplementary Information for "A decision-making tool to navigate through Extracellular Vesicle research and product development"

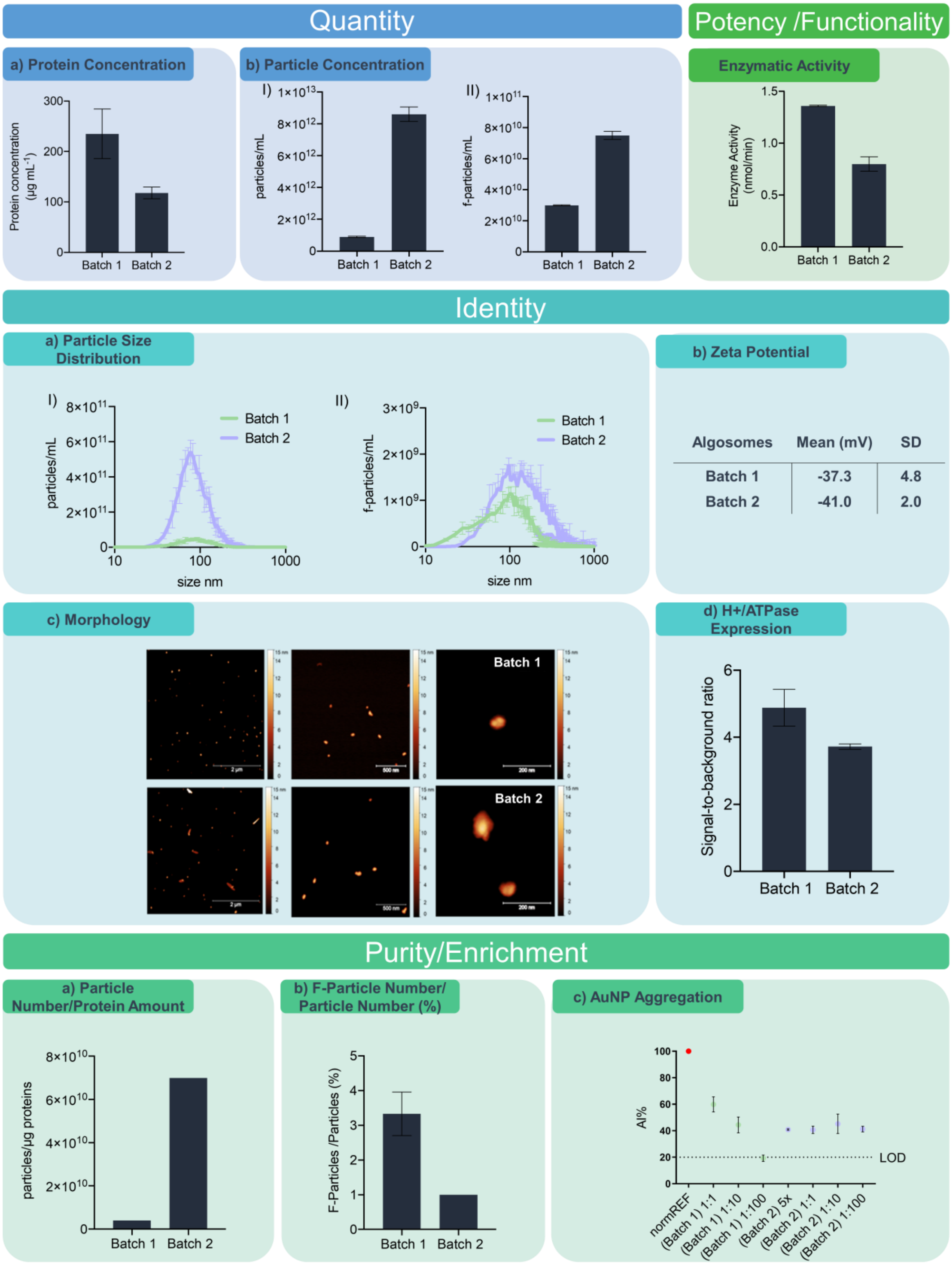

**Figure S1. Quantity**: a) Protein quantification performed on Batch 1 and Batch 2. Protein concentrations were measured as previously described [18]. Data are expressed as average µg/mL ± standard deviation of two technical replicates. b) Concentration measurements of scattering nanoparticles (I) and Di-8-ANEPPS-labelled fluorescent nanoparticles (II) of Batch 1 and Batch 2. Analysis was performed as previously described [18]. **Potency/Functionality**: Esterase activity assay performed on Batch 1 and Batch 2. Esterase activity was assessed by a recently patented Esterase Activity Assay [25,26]. Data are expressed as average nmol/min ± standard deviation of two technical replicates. **Identity**: a) Size distributions of scattering nanoparticles (I) and Di-8-ANEPPS-labelled fluorescent nanoparticles (II) assessed on Batch 1 and Batch 2. Analysis was executed as previously described [18]. b) Z-potential distribution profiles of Batch 1 and Batch 2. Table displays mean and standard deviation of Z-potential values, expressed in mV, resulting from three technical measurements. Z-potential was assessed using the ZetaView PMX-120 instrument by Particle Metrix (Germany), which is equipped with a 488 nm laser (40 mW power) and a CMOS camera sensor. Instrument initialization and camera alignment were executed according to the manufacturer's guidelines, utilizing 100 nm polystyrene size standard beads. Measurements were performed in continuous mode, diluting samples with a buffer having a conductivity equal to or below 2000 µS/cm, namely 0.1X concentrated PBS solution; data are expressed in millivolts (mV), along with the average percentage of frequency based on three technical measurements. c) In-air AFM-based morphology analysis performed on Batch 1 and Batch 2. Analysis was carried out as previously described [18]. d) H+/ATPase expression in Batch 1 and Batch 2. Analysis was performed by direct ELISA. Briefly, each sample was loaded in duplicate onto a transparent and uncoated 96-well microplate with high-binding capacity (HansaBioMed Life Sciences) for overnight nanoalgosome capture at 37°C. After three washing steps with 300 µL/well of washing buffer (HansaBioMed Life Sciences), microplate wells wells blocked with 300 µL/well of sample buffer (HansaBioMed Life Sciences) for 2 hours at room temperature (RT). Following a second washing step performed as described above, microplate wells were incubated with primary rabbit antibodies anti-plasma membrane H+ATPase (Agrisera; Cat.: AS07 260; 1/1000 dilution in sample buffer) for 2 hours at 37°C. A subsequent third washing step performed as aforementioned was followed by a microplate well incubation with Horse Radish Peroxidase (HRP)-conjugate secondary antibodies (HansaBioMed Life Sciences; 1/5000 dilution in sample buffer) for 1 hour at 37°C. The final procedural steps were performed according to ExoTEST™ manufacturer´s protocol (HansaBioMed Life Sciences). ODs were measured at 450 nm using Tecan GENios Pro microplate reader. Data are expressed as average signal-to-background ratio ± standard deviation of two technical replicates. **Purity/Enrichment**: a) Yields of Batch 1 and Batch 2 in terms of nanoalgosome protein content and particle number. Data were obtained as previously described [27]. b) Purity of Batch 1 and Batch 2 in terms of fluorescent particles over total particles. Analysis was based on the results obtained from the assessment conducted using the NanoSight NS300 instrument, as above reported, and applying the formula (f-particles/total particles)X100. c) CONAN assay performed on Batch 1 and Batch 2. Assessment was performed as previously described [23]. Data are expressed as mean ± standard deviation of three technical replicates. Legend: normREF = pristine gold nanoparticles (AuNPs) used for normalization; LOD = assay threshold for the detection of co-isolated proteins (Aggregation Index or AI% = 20%).

| Assessment Criteria | Assessment Sub-Criteria | Importance Weights (W) | Alternative X (Nanoalgosome Batch X) | |
|---|---|---|---|---|
| | | | Analytical Assessment Score (AA) | Assessment Criterion Score (AAxW) |
| **Quantity** | Particle number per volume unit | 3 | 3 | 9 |
| **Identity** | Particle size distribution | 2 | 3 | 6 |
| | Morphology | | 3 | 6 |
| | Membrane protein H+/ATPase expression | | 3 | 6 |
| | Surface charge | | 3 | 6 |
| **Purity/Enrichment** | Particle number-to protein amount ratio | 3 | 3 | 9 |
| | Percentage of total fluorescently-labelled particle number/total scattering particle number | | 3 | 9 |
| | AuNP aggregation | | 3 | 9 |
| **Potency/Functionality** | Enzymatic activity | 2 | 3 | 6 |
| **Actual Final Grade** | | | | 66 |
| **Actual Final Grade/ Theoretical Final Grade (%)** | | | | 100 |

**Table S1.** Calculation of theoretical final grade. Legend: X = unknown nanoalgosome batch.
